# Frequently Together, Among Model Systems Molecular Alterations: Cancer Maintenance Markers with Translational Study Design

**DOI:** 10.1101/2024.01.26.577415

**Authors:** Maya Ylagan, Qi Xu, Jeanne Kowalski

## Abstract

In contrast to cancer driver gene alterations, genes with alterations occurring frequently together are unlikely to play a direct role in tumor development, and more likely to contribute to normal cellular functions necessary for the ongoing maintenance and survival of the tumor, with the most robust of these genes preserved across model systems. We introduce two molecular signature-centric methods, RECO (recurring and co-occurring) and Crosstalk, to identify gene sets characterized by molecular alterations that occur frequently together across sample model systems. Our overall approach builds a cancer information exchange among samples systems based on an analysis of gene and sample-level signature pairs using RECO and Crosstalk methods. The results may be used to design translational studies from gene signature discovery in patients to its application in cell lines and patient-derived xenograft models, alongside the discovery of potential cancer maintenance markers. We demonstrate the capabilities of our approach by exploring the discovery of frequently together molecular alterations between patient tumors and cell lines, and between histologically similar tumors with different sites of origin, and their associations with outcomes in several cancers. As understudied, expanded markers, genes with frequently occurring molecular alterations between systems offer potentially new insights into similar cancer etiology and treatment targets outside the norm, in addition to informing on translational study design.

**Availability and implementation:** The method is implemented at: https://github.com/kmlabdms/CIERCE

## Introduction

Cancer remains a leading cause of morbidity and mortality worldwide, with its complex and heterogeneous nature, posing ongoing significant challenges for diagnosis, treatment, and prevention. Oncogenes and tumor suppressors are well known markers for prognosis, notably for their effects on cancer related pathways, and they have been studied since the discovery of SRC in 1911^1^ for their roles in diagnosis, cancer subtyping, and targeted treatment^2^. A mutation in an oncogene acts upon a molecular pathway as a single actor, promoting or removing hinderances for oncogenesis. This oncogene paradigm finds one gene to be at the root of the disease^3^. For instance, metastatic melanoma with the BRAF V600E mutation has a known pathway that it impacts and there is a known mechanism for carcinogenesis. With a known biological mechanism, pharmaceuticals can be developed to target that abnormality, and in this case, vemurafenib^4^. For each unique biological cause of cancer, it is unreasonable to think that we can find a mechanism and a targeted therapy. Notably, not all occurrences of cancer originate from oncogenes and tumor suppressors, such as the case of rare cancers, and making it challenging to find targeted therapies for these cases. While studies into oncogenes have undoubtedly advanced our understanding of specific genetic aberrations that drive malignant transformation, they inherently present an incomplete picture of the intricate nature of interactions governing cancer.

Moving beyond the confines of the oncogene paradigm, cancer is a complex and dynamic system biology disease, created by multistep processes. In the intricate and diverse cancer biological system, herein, we aim to identify a set of genes acting as dependencies in cancers that are largely outside of oncogenes. These genes may not necessarily play a direct role in tumor development but rather contribute to normal cellular functions necessary for the ongoing maintenance and survival of the tumor^5^. We posit such maintenance gene markers to occur often and together among sample model systems, such as between patient tumors and patient-derived-cell lines and xenografts (PDX). This definition is an extension of the oncogene paradigm to acknowledge the dependencies of a tumor more generally in cancer research.

The overall goal of identifying molecular alterations that occur often and together among samples from different model systems aims to unravel gene sets that are likely to reflect putative dependencies in cancers that may pose as shared treatment targets and markers of common outcomes. To address this goal, we present CIERCE (Cancer Information Exchange for Recurring, Co-occurring, and Co-Enriched genes), as a sample-signature-centric method in which to generate, explore, and relate molecular signatures within and among multiple systems. As a method, CIERCE contains two processes: RECO (Recurring and Co-occurring) and Crosstalk. To address the interconnected nature of cancer dependent genes, we define the molecular alterations among gene groups that occur often and together as REcurring and CO-occuring (RECO) within a system. Using sample-level gene signatures, the most recurring genes across all samples are clustered based on their within-sample prevalence to find groups of co-occurring genes, producing RECO gene groups within a system.

RECO gene groups can also be found between model systems, such as among patient tumor samples, cell lines, and patient-derived xenografts and organoids. To find RECO gene groups present in multiple systems, we introduce a Crosstalk analysis that is structured around the hypergeometric distribution. Crosstalk employs a sample-pairwise analysis to find cross-system, co-enriched genes, i.e. Crosstalk genes. Using Crosstalk genes, RECO gene groups can then be found among multiple systems. This novel functionality facilitates the exchange of molecular signature information and insights between samples and systems, revealing commonalities and interactions with potential therapeutic and clinical outcome implications.

As a method, CIERCE enables efficient, whole-systems experimental design by making use of information across model systems. As example, in a ‘bench-to-bedside’ experiments, results from an *in vitro* cell line experiment are interrogated using public resource, patient-tumor-derived data that if clinically impactful, a subsequent search for PDX/PDO models is performed to design an in vivo experiment. In this setting, three model systems, cell lines, patient tumors, and PDX’s are interrogated at different stages of the experiment for the same signature. Alternatively, using a crosstalk analysis, the signature could be interrogated in all three systems for efficient design and a subsequent RECO analysis performed to identify sub-signature gene groups common among them and distinct to each system. Thus, CIERCE as a method takes this traditionally multi-step, somewhat disconnected design process and enables a cohesive, signature sample-level coenrichment analysis of prevalence within and among model systems. CIERCE also enables the discovery of expanded markers for cancer etiology and treatment. Besides oncogenes, there has been a paucity of gene markers that have been translated to the clinic. By defining frequently occurring together molecular alterations between model systems, researchers can explore an expanded universe of markers with yet untapped potential. Thus, CIERCE offers researchers the opportunity to uncover new connections and biological mechanisms underlying cancer progression.

With the advancements in high-throughput genomic sequencing, there has been generation of large volumes of publicly available cancer data^6,7^. The CIERCE method is applicable to any signature defined at a gene-level and any samples for which gene signatures may be defined. Thus, there is no restriction to molecular data type or sample types. We provide an R package with functions to run both RECO and Crosstalk analyses to derive gene sets that are often, together, and between systems as potential multi-gene markers of cancer etiology and treatment. We demonstrate the method’s utility of frequently together, between systems, molecular gene signatures by applying CIERCE to investigate cancer sample-level molecular signatures defined by expressed gene amplifications and mutations within and among model systems based on common histology (Case 1, 3), underlying biology (Case 4), and translational bench-to-bedside study design (Case 2).

## Results

### Architecture of CIERCE

CIERCE focuses on the analysis of sample-level signatures. The multi-omics analysis methods we present within CIERCE — RECO and Crosstalk — can be used in two different analysis workflows depending on the number of systems. For the workflow within a single system, all sample signatures can be used in the RECO analysis method to define sets of RECO genes groups within that system of samples (**Fig. 1a**). For the workflow for two (or more) systems, sample-level signature pairs are used in Crosstalk to define co-enriched signatures between them (**Fig. 1b**). Co-enriched signature genes can be used in a subsequent RECO analysis to detect frequently together, between systems, signature gene groups (**Fig. 1c**). These two analysis workflows offer insights into the broader implications and commonalities of signatures between or within system(s).

**Figure 1.**
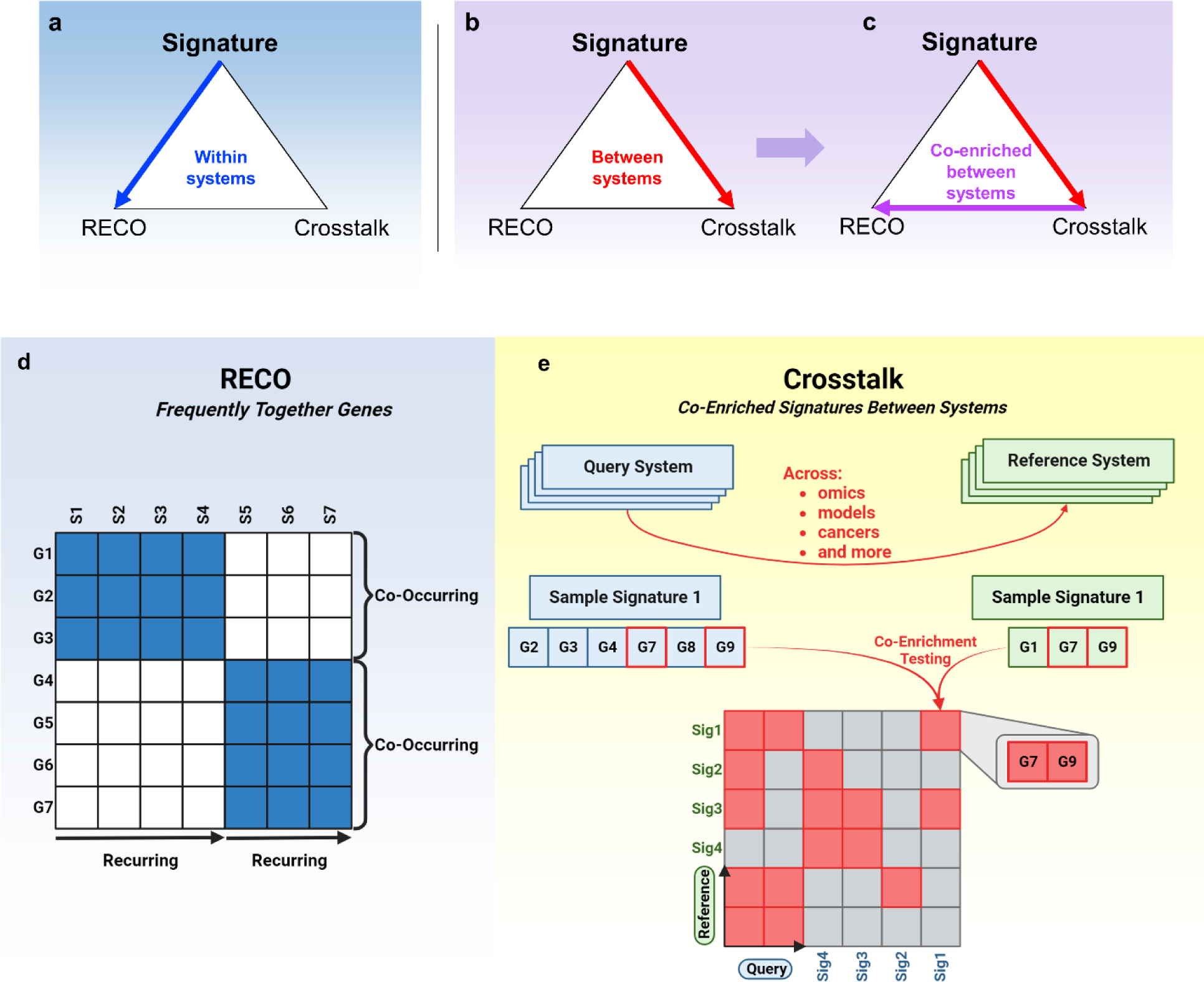
An analysis of recurring and co-occurring gene prevalence combined with the assessment of sample pair co-enriched gene signatures define the build of a Cancer Information Exchange of Recurring, Co-occurring, and Co-enriched Gene Signatures (CIERCE). **a**, Signature to RECO. The within-system analysis defines recurring and co-occurring (RECO) gene groups based on a prevalence clustering of gene signatures. **b**, Signature to Crosstalk. This between-system analysis defines a Crosstalk between systems (e.g., cancer types, data sources, patient-derived models) that is based on significantly co-enriched genes signatures among systems. **c**, The between-system co-enriched genes from Crosstalk can be further used to define RECO gene groups. **d**, The introduction of the RECO process to define frequently together gene signature groups. RECO genes are recurring across samples (often) and co-occur within samples (together). Columns are samples (S1-S7), and rows are genes (G1-G7). A visual grouping of co-occurring and recurring are provided in reference to the clustering of the samples and genes. **e**, The introduction of the Crosstalk process to define co-enriched gene signatures among systems. Crosstalk defines genes that are significantly co-enriched between defined query-reference pairs to form a distinct signature gene set that is co-enriched across systems. For the example systems, a single signature from each system is shown to illustrate the framework for pairwise co-enrichment testing. The co-enriched genes are stored in a matrix of query-reference pairs.

Crosstalk uses sample-to-sample pairwise signature enrichment between two systems’ signatures to uncover meaningful connections and interactions from their co-enriched genes between systems (**Fig. 1e**). These systems can be any set of gene signatures, most commonly a biological phenotype or genotype for any organism; for instance, amplification signature sets, cancer biomarker gene sets, or pathways of interest can all used as systems for Crosstalk analysis. Crosstalk is an adaptation of an enrichment analysis based on the hypergeometric distribution and uses two groups of gene sets, one of which is defined as a query and the other, a reference gene set. In this setting, the idea is to interrogate query gene sets for enrichment of reference gene sets. As an example, the typical setting of identifying enrichment of a given signature such as DNA homologous recombination (HR) in a TCGA-derived molecular subtype signature, herein, HR would be the reference signature that is queried in the subtype signature. As a further example, to identify the enrichment of TCGA-derived sample-level molecular signatures in CCLE-derived cancer cell line-level molecular signatures, the former would be defined as the reference and the latter, the query gene signature sets.

A crosstalk analysis identifies, for each query-reference gene set pair, the enrichment of the query gene set in the reference gene set in relation to the background gene pool. Specifically, we define two different model systems such that *I* and *J* refer to the reference and query sample identifiers respectively. We further define *P* and *C* such that and where each i^th^ and j^th^ sample of *P* and *C* respectively contains a set of signature genes. Denoted by the following:

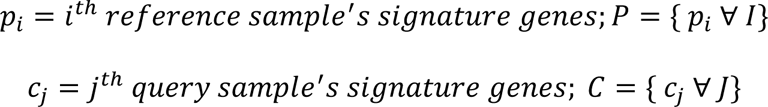

Define an *I* × *J* crosstalk matrix, *G* ∈ ℝ^2^ where each entry, *G*_*i*,*j*_ = *p*_*i*_ ∩ *c*_*j*_ *iff Pr*[*p*_*i*_ ⊆ *c*_*j*_| c_*j*_ ∀ *J*] ≤ 0.05 using the hypergeometric distribution. We opted for a hypergeometric test of enrichment over testing for overlap to consider the different model system backgrounds from which signatures are derived. Using the resulting matrix *G* of crosstalk genes, the set of distinct crosstalk genes, *g*, is the union of each element of *G*.

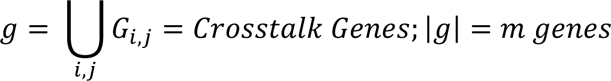

Crosstalk results in finding *g*, the set of genes whose sample query signatures are significantly enriched in a reference sample among all reference-query sample pairs. Both query and reference gene sets may be defined according to biological questions and analyzed using various combinational patterns.

The RECO algorithm produces RECO gene groups which are a set of genes that occur together and often. RECO gene groups are defined as recurring (RE) and co-occurring (CO) signature genes across samples. Recurrence is defined by a user-determined level of prevalence of signature genes among samples. Co-occurrence is defined by the presence of two or more signature genes among samples. To find co-occurring signature genes, row-wise unsupervised clustering on the recurring signature genes is performed based on their prevalence, binary representation as present in samples (**Fig. 1d**). CIERCE identifies the resulting clusters as RECO gene groups, revealing potential interactions and relationships among the genes that define them. More specifically, for a set of genes, *g* (either from Crosstalk or not), with cardinality *g*. We present the following method to find partitions of *g* such that sample prevalences are similar. We define a set of *n* samples’ signature, *S*, whose signatures are filtered to be a subset of *g*.

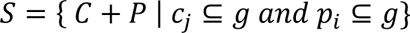

These samples are used to compute the *m* × *n* binary matrix, *B*. With elements defined by the indicator function, II, which denotes if the sample signature *S*_*n*_ contains the *m*^th^ gene.

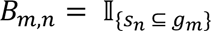

The resulting matrix *B* is then subjected to row-wise k-means clustering to find the recurring and co-occurring gene signatures between model systems. To interrogate the prevalence of this RECO gene group in the population of samples, individual samples must be determined to contain the entire RECO gene group.

### A Pan-Can EAS Analysis using CIERCE

We explore the prevalence of expressed and amplified signatures (EASs) within pan-cancer (Pan-Can) analyses. EAS genes are included in the sample’s signature if the gene has high CN and high RNA expression. By integrating these data types into a single signature, CIERCE identifies genes that exhibit both amplification and expression at the sample-level, allowing for a comprehensive comparison of the functional consequences of gene amplification and expression between systems and samples.

To create EASs for Crosstalk and RECO analyses, all available TCGA data was used to analyze EASs across cancer type; this encompassed a total of 8,651 out of 9,892 TCGA samples with available expression and amplification data to contain an EAS, and their corresponding EASs have been populated into CIERCE for others to run their own analyses on. Included in the EASs are genes that are present in samples containing at least one EAS gene. EASs are represented in greater than 95% of samples from 17 different cancer types, including notably, in BRCA with the most overall EASs (**Supplemental Fig. 1a**). The distribution of the number of genes for each EAS varies among cancer type, and there is no clear relationship between the number of samples with EAS and the median number of genes in an EAS (**Supplemental Fig. 1b**). We further investigated the overall survival (OS) outcomes associated with containing an EAS across multiple cancer types (**Supplemental Fig. 1c**). The following Pan-Can project’s samples with EASs were significantly associated with shorter OS compared to samples without an EAS: LGG (HR= 2.7, CI= [1.2,6.2]) and UCEC (HR= 2.5, CI= [1.2,5.0]). By comparison, CESC (HR= 0.1, CI= [0.02,0.4]) EAS signatures were significantly associated with a favorable, longer OS. Altogether, our Pan-Can analyses sheds light on the landscape of EASs in terms of their prevalence in various cancer types that are highly represented and in terms of their survival associations, both show support for the further study of their potential role as biomarkers.

EAS creation and landscape analysis was also repeated for CCLE cell lines. EASs were found for 891 cell lines of the 923 that had the applicable data available. CCLE cell lines had a majority of primary sites having a very high prevalence of EAS (**Supplemental Fig. 2a**). The distribution of the number of genes for each EAS varied by primary site and did not show a clear relationship to the number of samples with EAS and the median number of genes in an EAS (**Supplemental Fig. 2b**).

### Case study 1: RECO analysis of EASs in low grade glioma (LGG)

Considering the significantly shorter OS in the LGG TCGA project associated with having an EAS compared to samples without EAS (**Supplemental Fig. 1c**). As a case study, we explore RECO gene groups in this cancer type using the workflow outlined in **Fig. 1a**. To obtain RECO gene groups, clustering on binary prevalence of the most recurring EAS genes (n=28) was done (**Fig. 2a**). The RECO gene groups (n=6) are shown in an oncoplot to reflect their prevalence across samples (n=511). The chromosomal location of these genes was assessed and RECO gene groups were found to be interspersed with each other rather than isolated (**Supplemental Fig. 3a**). These genes are all on chromosome 7, and among them, *SMO* is an oncogene^8,9^. To find the prevalence of the RECO gene groups within a sample, the sample must have all genes in the RECO gene group present. For each RECO gene group, their population-wise prevalence in samples was found and their IDH1 mutation status was visualized (**Fig. 2b**). An additional feature of this clustering is that it can be run on not only binary prevalence, but also values. In **Fig. 2c**, we use copy number values to cluster the most prevalent genes demonstrating the difference to **Fig. 2a**’s RECO binary prevalence clustering output allowing users to find dose-dependent gene groups.

**Figure 2.**
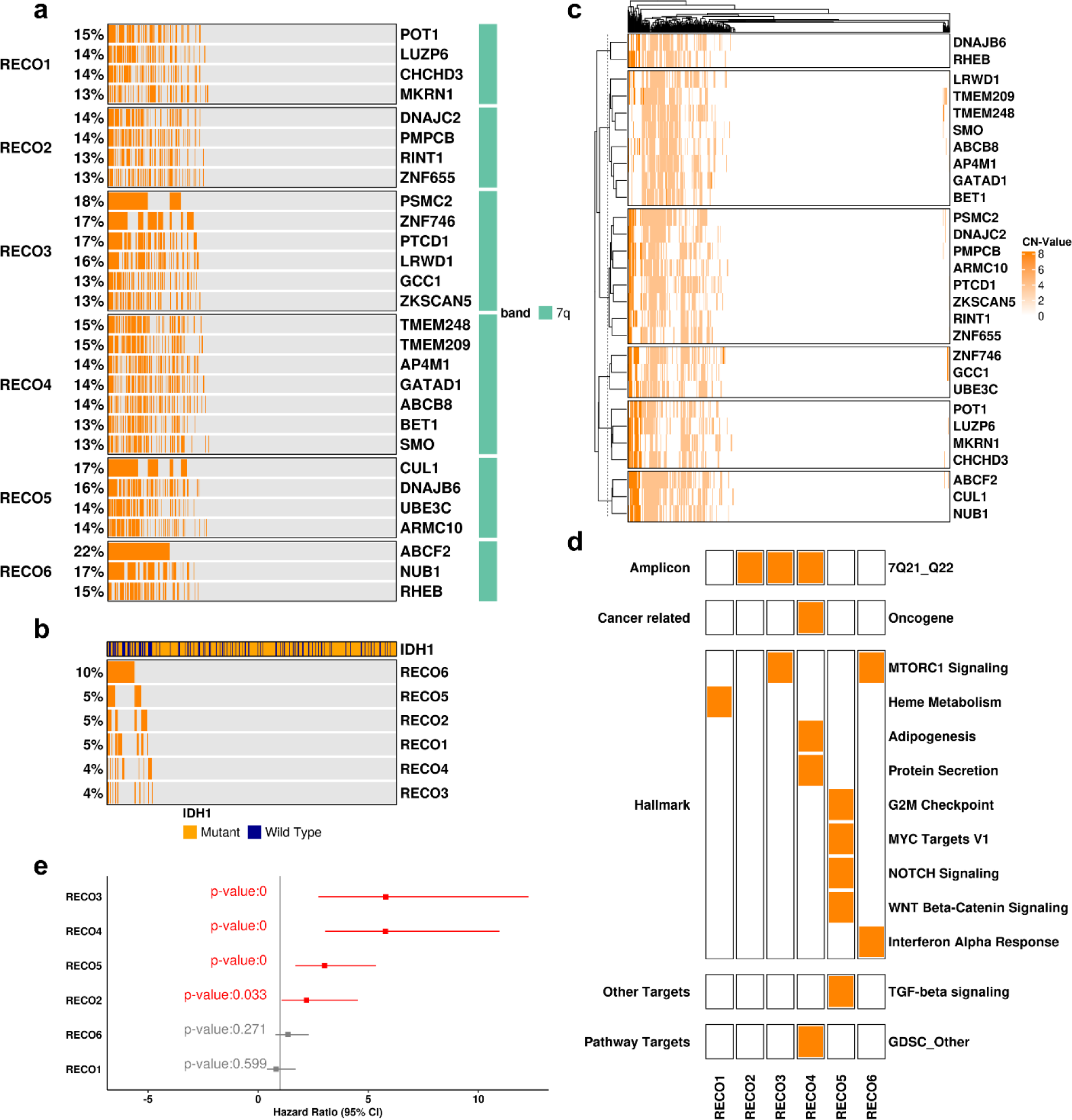
A RECO Analysis of sample-, gene-level expressed amplification signatures (EAS) within TCGA Low Grade Glioma (LGG) patients reveals several non-oncogene-containing gene sets, except for one, associated with significantly shorter survival outcomes. **a**, An oncoplot of top individual EAS gene prevalence clustering results that define RECO gene cluster groups in TCGA LGG. **b**, A summary prevalence oncoplot of samples that contain all genes within RECO gene group clusters. **c**, Gene-level copy number value clustering showing the most prevalent copy number altered genes in TCGA LGG within RECO gene group clusters. **d**, Crosstalk enrichment of RECO gene groups as the query against known signatures as reference. Significant enrichments (p<0.05) are indicated in orange. **e**, Forest plot of hazard ratios (HR) and 95% confidence intervals for overall survival based on comparing samples within each RECO gene set versus not in the set. Significant HR’s are highlighted.

To gain a more functional understanding of the RECO gene groups, we employ Crosstalk as an enrichment testing framework using the RECO gene groups as signatures and following the workflow depicted in **Fig. 1b**. With defined RECO gene groups in **Fig. 2a**, a Crosstalk using known signatures as the reference and the RECO gene groups as the query was done to show the enrichment of the RECO gene groups in known signatures (**Fig. 2d**). From using Crosstalk as an enrichment testing framework, the enrichment of multiple Hallmark signatures in the RECO gene sets was found. Expanding upon the functional exploration, co-expression networks were found for each RECO gene group to find genes that were commonly expressed across the interrogated TCGA samples, i.e. hub genes, in yellow (**Supplemental Fig. 3b**). Further assessing the RECO gene groups CIERCE produced, within each RECO gene group an association test using Fisher’s exact test was done to compare the association of IDH1 mutations with each RECO gene group, and RECO1 was the only insignificant association while for RECO 2, 3, 4, 5, and 6 all had a significant association (p<0.001) compared to RECO prevalent samples (**Supplemental Table 1**). Significant associations were also found for Fraction Genome Altered (FGA) with RECO1 having the only insignificant association while for RECO 2, 3, 4, 5, and 6 all had a significant association (p<0.001) compared to RECO prevalent samples (**Supplemental Table 1**). We further investigated the overall survival (OS) outcomes associated with the prevalence of each RECO gene group. RECO3 (HR= 5.80, 95% CI = [2.73,12.28]), RECO4 (HR = 5.77, 95% CI = [3.04,10.95]), RECO5 (HR = 3.00, 95% CI = [1.68,5.35]), and RECO2 (HR = 2.20, 95% CI = [1.06,4.52]) were significantly associated with shorter OS compared to samples without the given RECO gene group (**Fig. 2e**).

### Case study 2: Breast cancer patient tumors and cell lines crosstalk analysis

To compare model systems within a cancer type we focused on HER2+ breast cancer using the workflow outlined in **Fig. 1b, c**. Using TCGA BRCA HER2+ samples as the query (n=159) and CCLE HER2+ as the reference (n=51) a Crosstalk was run (**Fig. 3a**). This Crosstalk was further visualized with a 3D heatmap to accentuate the patient and cell line pairs with high gene overlap (**Fig. 3b**). There were 4,021 genes resulting from Crosstalk. To assess the sample size considerations between systems, we interrogate the signature genes used in Crosstalk. In Crosstalk for a given gene, how many samples contain that gene in the query compared to the reference by computing the residual to y=x for that gene (**Supplemental Fig. 4a**).

**Figure 3.**
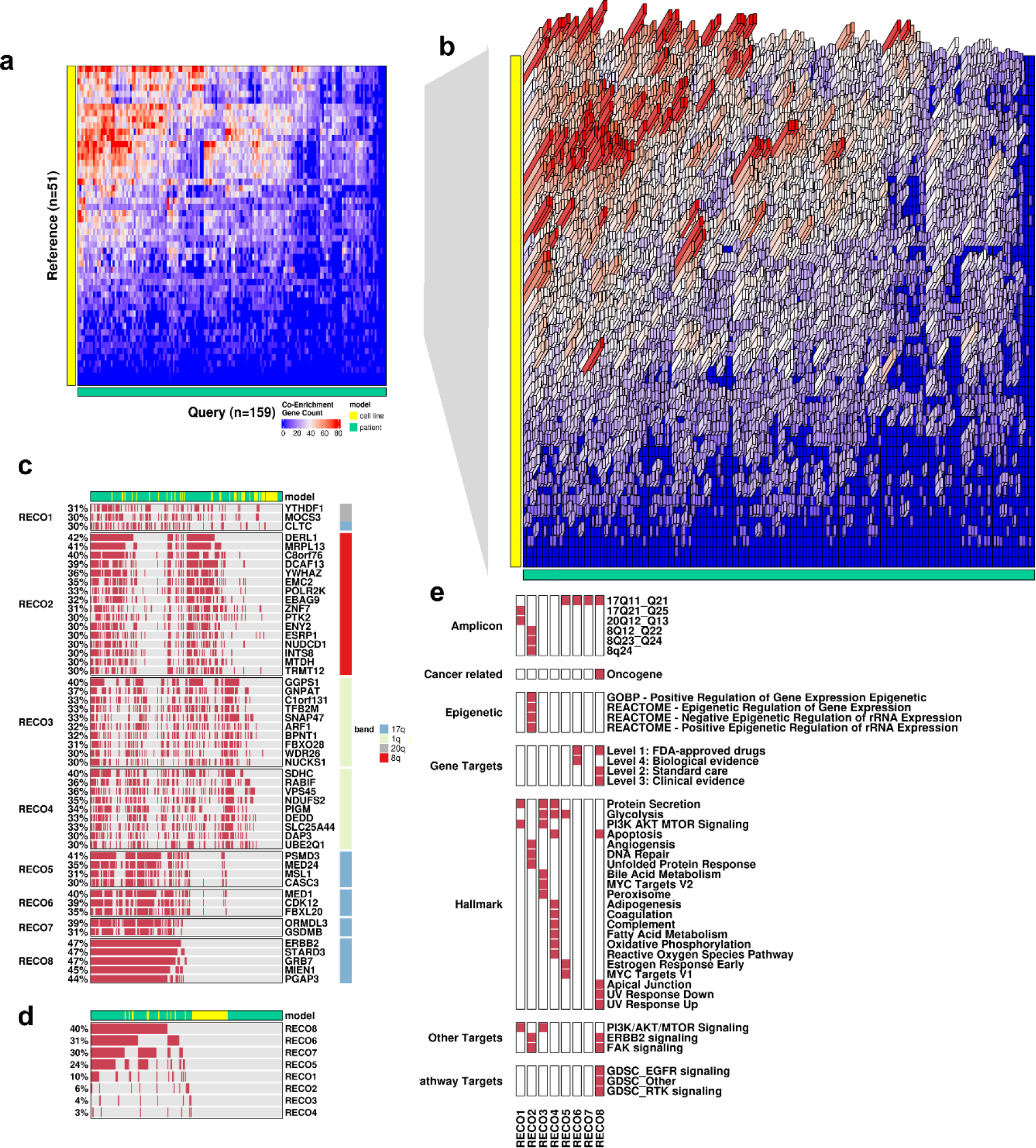
A crosstalk analysis of sample-, gene-level expressed amplification gene signatures (EAS) between HER2+ TCGA breast cancer patients and HER2+ CCLE breast cancer cell lines reveals several common non-oncogene-containing gene sets, except for one. **a**, Crosstalk of HER2+ TCGA breast cancer patients EAS’s as Query samples (n = 159) against HER2+ CCLE breast cancer cell lines EAS’s as reference (n=51). Heatmap of the crosstalk 159 x 51 matrix define the number of EAS gene signatures significantly co-enriched between patient-cell-line pairs. **b**, Three-dimensional heatmap of the Crosstalk matrix in **a** to visualize the pairs of samples with the highest co-enrichment. **c**, Oncoplot visualization of RECO gene groups prevalence derived from the distinct list of significant co-enriched pairs EAS signatures analyzed by the Crosstalk process. **d**, Oncoplot visualization summary of the prevalence of HER2+ TCGA and CCLE cell lines breast cancer samples with EAS RECO gene signatures. **e**, Crosstalk enrichment of RECO gene groups from **c** as query against known signatures as reference. Significant enrichments (p<0.05) are highlighted. **f**, Residual plot assessing the sample size considerations between systems by interrogate the input genes into Crosstalk. For a given gene the number of samples contain that gene in the query compared to the reference by computing the residual to y=x for that gene.

After running Crosstalk, RECO was run on the resulting genes from Crosstalk for both patient and cell line samples to find RECO gene groups (n=9) for the most prevalent genes (n=52) in common to both (**Fig. 4c**). *ERBB2* (HER2) was unsurprisingly found in a RECO gene group as all samples were HER2+; interestingly only 47% of these HER2+ samples were prevalent for this EAS gene which was clustered with other genes on chromosome 17q, illustrating the power of RECO to help inform and design wet lab experiments for possible associated targets. Clinical associations within each RECO gene group across samples that were either prevalent or not for that RECO gene group were done finding that there were multiple demographic associations with RECO gene groups (**Supplemental Table 2**). The chromosomal location of these genes was assessed (**Supplemental Fig. 5a**) to investigate spatial relationships between RECO gene groups, demonstrating the RECO groups were often isolated rather than interspersed. These RECO gene groups were then interrogated for prevalence in samples (**Fig. 3d**) and like in the prior case study used in a Crosstalk with known signatures to find enrichment in these known signatures (**Fig. 3e**). When testing prevalence of a RECO gene group as an indicator for survival no RECO gene groups showed any significant effects. Expanding upon the functional exploration, co-expression networks were found for each RECO gene group to find genes that were commonly expressed across the interrogated TCGA samples, i.e. hub genes, in yellow (**Supplemental Fig. 5b**)

**Figure 4.**
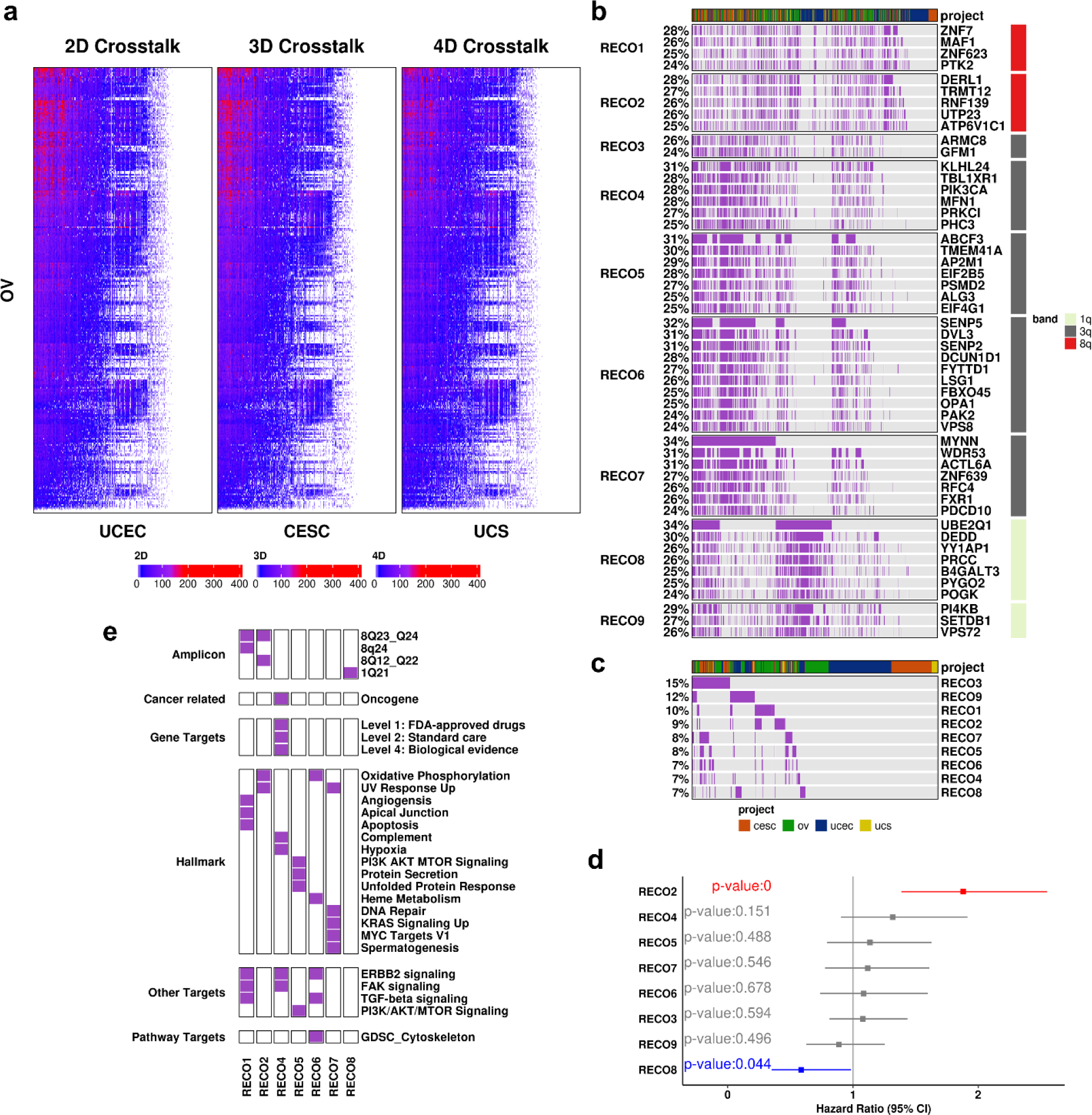
A nested crosstalk series analysis of sample-, gene-level expressed amplification gene signatures (EASs) among ovarian serous cystadenocarcinoma (OV), uterine corpus endometrial carcinoma (UCEC), cervical squamous cell carcinoma and endocervical adenocarcinoma (CESC), and uterine carcinosarcoma (UCS) cancer types that define a TCGA Pan Gynecological Cancer (TCGA Pan-Gyn-Can) cohort reveals several non-oncogene-containing gene sets, except for one, with significant favorable and unfavorable survival. **a**, Nested Crosstalks of sample-level EASs between TCGA OV patients as reference samples (n = 297) against TCGA UCEC (n=521), CESC (n=291), and UCS (n=56) as queries. The heatmap of the crosstalk matrices define the number of EAS genes significantly co-enriched sample pairs. **b**, Oncoplot visualization of RECO gene groups derived from the distinct list of significant co-enriched EAS gene signatures analyzed by the Nested Crosstalk analyses from **a**. **c**, Oncoplot visualization summary of the prevalence of RECO gene groups among all samples that define TCGA Pan-Gyn-Can cohort. **d**, Forest plot of hazard ratios (HR) and 95% confidence intervals for overall survival based on comparing samples containing RECO EAS genes versus not. Models that violated the proportional hazards assumption were removed. Significant HR’s are highlighted. **e**, Crosstalk enrichment of RECO gene groups from **b** as query against known signatures as reference. Significant enrichments (p<0.05) are highlighted.

### Case study 3: Pan-Gynecological-Can (Pan-Gyn-Can) EAS crosstalk analysis

To illustrate the capabilities of Crosstalk in exploring and uncovering EAS genes across different cancer types, we present a case study focusing on gynecological cancer due to significant survival differences found in UCEC and CESE (**Supplemental Fig. 1c**) and the published use of similar cohorts ^10^.

To find commonalities between these Pan-Gynecologic-Can (Pan-Gyn-Can) cancer types we must use Crosstalk to compare more than one system using the workflow outlined in **Fig. 1b, c**. Here we demonstrate how to use Crosstalk with greater than 2 systems; we use a nested approach. If a nested approach is not used, then the results will not be representative of all the systems. The reference will be represented, but the samples in the query systems will not be tested for co-enrichment amongst themselves. This Nested Crosstalk uses 2 systems as a baseline reference and query, their sample pairs are used as a reference for further query systems. For a more concrete explanation, the Pan-Gyn-Can Nested Crosstalk was done first using a reference as TCGA OV against a query of TCGA UCEC, the resulting sample pairs were used as a reference and CESC was used as query. The resulting sample pairs are kept in relation to the original OV and UCEC but the co-enriched signature is now representative of OV, UCEC, and CESC; these pairs are used as reference and UCS used as a query (**Fig. 4a**). Using Nested Cross-talk on the Pan-Gyn-Can cohort produces common EAS genes between these Pan-Gyn-Can cancers. There are 10,054 unique co-enriched EAS genes in common between these Pan-Gyn-Can cancers. To assess the sample size considerations between systems, we interrogate for each Crosstalk for a given signature gene how many samples contain that gene in the query compared to the reference (**Supplemental Fig. 4b**). Overall, there was a uniform prevalence of genes in reference than query seen by residuals to y=x for the first Crosstalk, for the second and third due to using sample pairs as reference these numbers are inflated and not informative nor interpretable but are presented for the reader to follow the methodology of the nested approach.

The Crosstalk genes were used in a RECO analysis to investigate the RECO gene groups across these two systems. **Figure 4b** represents the RECO analysis, where we detected top recurring genes (n=51) and 9 RECO gene groups. The sample level prevalence of these RECO gene groups were then calculated and visualized (**Fig. 4c**). Interestingly, an oncogene, PIK3CA, was in the top recurring genes^9,11^. Clinical associations within each RECO gene group across samples that were either prevalent or not for that RECO gene group were done finding a race association with RECO1 (p<0.05); FGA was significantly associated (p<0.05) with all RECO gene groups, and TMB was significantly associated (p<0.05) with all RECO gene groups except RECO9 (**Supplemental Table 3**). To investigate the spatial relationship of these genes, the chromosomal band location for these genes was visualized (**Supplemental Fig. 6a**) demonstrating these RECO gene groups are mostly spatially isolated with minimal mixing. To investigate the implications of these RECO gene groups on survival, an OS Cox model was run on each RECO gene group for the binary prevalence of the sample containing the RECO gene group; containing RECO2 (HR = 1.88, CI = [1.39,2.55]) and RECO8 (HR = 0.58, CI = [0.36,0.98]) had significant survival differences (**Fig. 4d**). To investigate the functional implications of these RECO gene groups, with the RECO gene groups as queries a Crosstalk was done against known signatures as reference to find the enrichment of these RECO gene groups (**Fig. 4e**). To note, the RECO gene group enrichment found multiple Hallmark signatures significantly enriched in RECO gene groups, allowing insight into these gene groups’ functionality. Continuing the functional analysis, co-expression networks for the TCGA samples were found for each RECO gene group to find genes that were commonly expressed across the interrogated TCGA samples, i.e. hub genes, in yellow (**Supplemental Fig. 6b**). This comprehensive analysis provides valuable insights into the functional impact of these EAS genes in gynecological cancers.

### Case study 4: Mutation signature crosstalk analyses between colorectal and pancreatic cancers

Pancreatic and Colorectal cancer share common genomic features, including presence of KRAS mutations^12^, germline mutations in BRCA1 and BRCA2^13^, and genetic alterations governing cancer progression, such as gene expression changes, copy number aberrations, chromosomal rearrangements, and epigenetic alterations^14^.

To demonstrate the capabilities of Crosstalk in exploring the relationship between other types of signatures, we present a case study focused on mutation signatures following the workflow from **Fig. 1b, c**. This analysis involves investigating the crosstalk between mutations in TCGA patient samples with mutational data present in colorectal (n=528) and pancreatic (n=176) cancer and the DNA Damage Response (DDR) gene set. Mutational signatures were defined as any mutated gene for that sample. Using a Nested Crosstalk approach as priorly described, Colorectal served as the reference, while pancreatic was used as the first query for Crosstalk, and DDR signature genes used as the second query. The study revealed 146 crosstalk genes common to both models and the DDR gene set, suggesting significant DDR activity cancer types’ mutational signatures (**Fig. 5a**). To assess the sample size considerations between systems, we interrogate for each Crosstalk for a given signature gene how many samples contain that gene in the query compared to the reference (**Supplemental Fig. 4c**). Overall, there was greater prevalence of genes in reference than query seen by residuals to y=x for the first Crosstalk, for the second Crosstalk due to using sample pairs as reference these numbers are inflated and not informative nor interpretable but are presented for the reader to follow the methodology of the nested approach.

**Figure 5.**
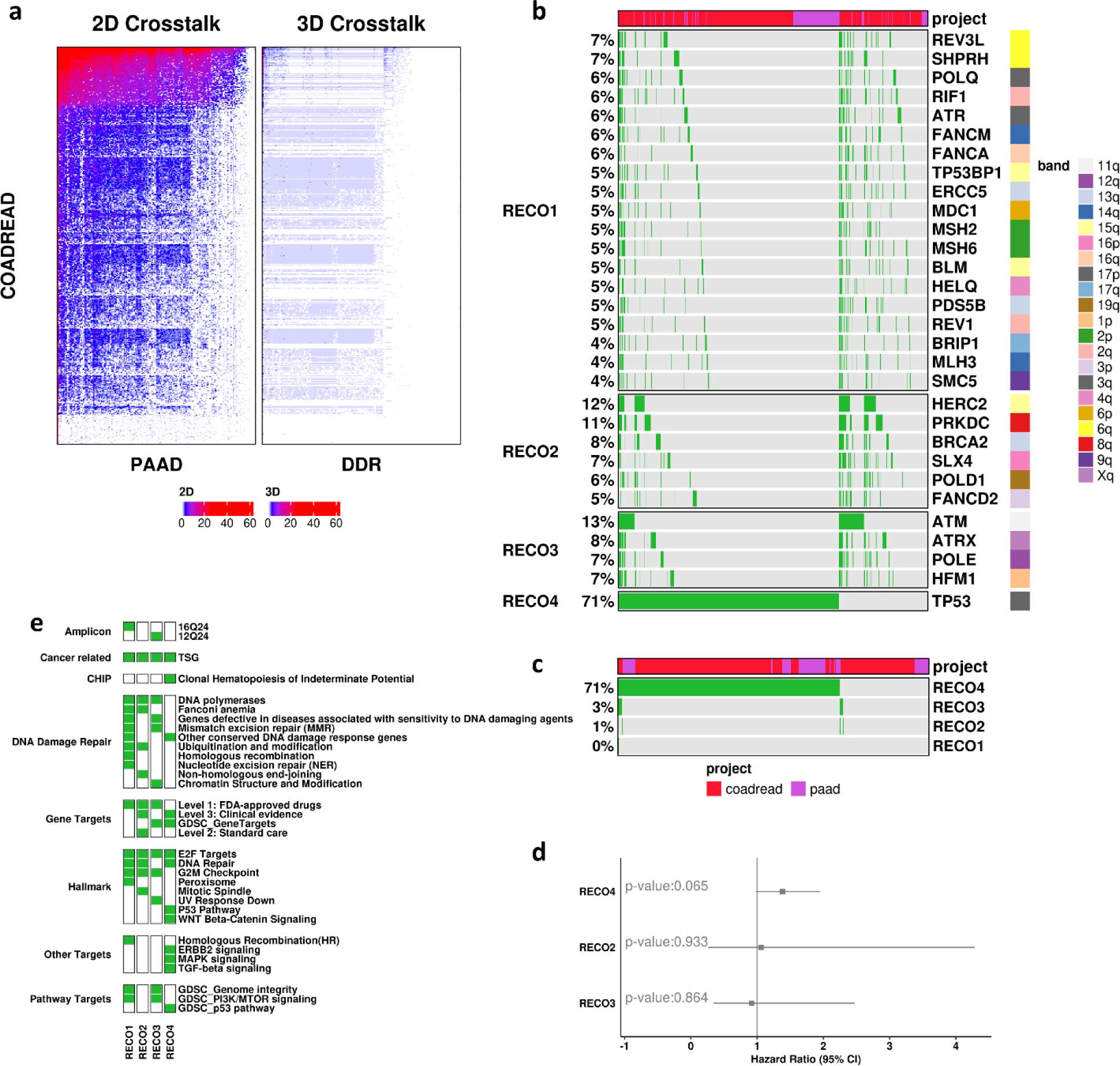
A nested crosstalk series analyses of sample-, gene-level mutation signatures among TCGA pancreatic and colorectal samples with DNA damage repair (DDR) gene signature demonstrate proof of principle based on the common prevalence of p53 in both cancer types that when mutated is associated with poor survival. **a**, Nested Crosstalk of sample gene mutation signatures between TCGA colorectal patients as reference samples (n = 528) against TCGA pancreatic patients as query (n=176), that is further tested against DDR signature genes (n=219 genes) as a second query. Heatmap of the crosstalk matrices define the number of genes in significantly co-enriched for each between system sample pair. **b**, Oncoplot visualization of RECO gene groups derived from the distinct list of significant co-enriched gene signatures analyzed by the Nested Crosstalk analyses from **a**. **c**, Oncoplot visualization summary of the prevalence of RECO gene groups among all TCGA pancreatic and colorectal samples. **d**, Forest plot of hazard ratios (HR) and 95% confidence intervals for overall survival based on comparing samples containing RECO gene mutations versus not. Models that violated the proportional hazards assumption were removed. Significant HR’s are highlighted. **e**, Crosstalk enrichment of RECO gene groups from **b** as query against known signatures as reference. Significant enrichments (p<0.05) are highlighted.

Further analysis detected four RECO groups for the most prevalent genes from Crosstalk (n=30) (**Fig. 5b**). The genes in RECO gene groups contain multiple tumor suppressors: MSH2, BLM, BRCA2, ATM, and TP53^9^. RECO4 contains a known tumor suppressor, TP53, as a singlet that does not co-occur with any other mutation as expected from a driver gene^9^. Clinical associations within each RECO gene group across samples that were either prevalent or not for that RECO gene group (**Supplemental Table 4,5,6**). Notably when assessing all patients and just coloreactal patients, TMB, FGA, and Stage were associated with RECO4 (p<0.05). When using all patients, multiple demographic associations with RECO gene groups were also found. To investigate the spatial relationship of these genes, the chromosomal location for these genes was visualized (**Supplemental Fig. 7a**) demonstrating the spread of these genes across the genome with no apparent connection to RECO gene groups. The high prevalence of RECO4 among all samples illustrates the shared genomic feature of these mutations in both cancer types (**Fig. 5c**). Samples prevalent with any RECO gene group were not found to have any survival impacts (**Fig. 5d**). To investigate the functional implications of these RECO gene groups, with the RECO gene groups as queries a Crosstalk was done against known signatures as reference to find the enrichment of these RECO gene groups (**Fig. 5e**) demonstrating that many DDR gene sets were found in the RECO gene groups along with other gene sets that could be associated targets in these cancers. Following the functional analysis, co-expression networks were found for each RECO gene group to find hub genes of expression (**Supplemental Fig. 7b**)

In summary, CIERCE provides researchers with a powerful method to explore the complex relationships between different gene signatures. By uncovering the connections and interactions between signatures and across systems, researchers can gain valuable insights into the underlying biology of various diseases.

## Discussion

CIERCE’s focus is to investigate how multiple genes act together in undefined ways. The RECO analysis introduced is a first of its kind method to find RECO gene groups in samples. Clustering has been a common theme for the analysis of omics data, the binary prevalence clustering that RECO employs enables the extraction of common sets of genes that occur throughout the set of samples. Other clustering analyses cluster all genes by quantized value, this enables a similar partitioning, but doesn’t provide a sample-centric perspective of the contents of the cluster. The RECO gene groups’ survival impacts demonstrate driver genes or cancer type are not sole indicators for disease and there is a need to investigate these gene groups. This demonstrates the need to be looking outside the scope of oncogenes and tumor suppressors, as the biological relationship between these genes in cancer is unknown; this methodology could guide the discovery of potential biomarkers and therapeutic targets.

A limitation of RECO is the parameters in which one sets. There is a tradeoff between the number of genes and the number of gene groups. The more this balance falls towards a larger number of genes in a RECO gene group the more negatively it affects the prevalence of the gene group signature across all samples, and vice versa. Depending on how the balance is set a gene singlet may appear, which is not a RECO gene group by definition.

Crosstalk is a first of its kind method to find enrichment of a system of samples in a system of samples. Other methods test a single sample/system of samples for enrichment in a gene or gene set, these methods are not comparable as they are testing enrichment in a completely different context^15^. Here we are finding co-enrichment of sample signatures rather than if a sample is enriched in a given external constant. Crosstalk in CIERCE enables the investigation of potential connections and interactions of signatures between different systems providing robustness of a finding across multiple model systems. Through the use of Crosstalk between systems, researchers can uncover new insights into the relationships between gene signatures and potentially identify common biological mechanisms underlying the compared systems.

Crosstalk is scalable to any other system(s) and numbers of systems. Regarding any other system, Crosstalk can not only find commonalities and compare across any genotype or phenotype, such as progression, grade, cancer type, and model system, but also amplification signature sets, cancer biomarker gene sets, and pathways of interest can all be used as systems for co-enrichment Crosstalk analysis. In general, Crosstalk can be used for any system of related samples, how this manifests in practice depends on the use case. Regarding any number of systems, Crosstalk can scale to any number of systems as demonstrated in the above case studies to allow researchers to design experiments more effectively and efficiently. The use of multiple systems creates both a more specific result due to the finer grained commonalities of the systems and a more robust result due to the shared significance of commonalities across the systems.

CIERCE methods – RECO and Crosstalk – can be used in many contexts. These methods can be vital for clinical providers to find public domain samples (i.e. TCGA or CCLE) that contain the same molecular alterations as their patients allowing for more personalized treatment regiments based on an increased knowledge base. This personalized medicine use case is not only for treatment options, but also for clinical trials. Primary samples that have been transformed into model systems can be analyzed with CIERCE methods to create inclusion criteria for the study. The applications of CIERCE methods are considerable and beneficial outside of cancer and across different biological disciplines. These methods enable a new perspective on current omics data for researchers to use as a study design tool to enable the finding of more comprehensive and robust targets for biological studies.

Future enhancements to CIERCE could include the integration of additional expression sources, such as from non-coding RNAs to further enhance our understanding of the levels of expression in relation to CN alterations. CIERCE can also be integrated with other analysis pipelines to serve as a standard downstream analysis from sequencing data. In terms of enhancing the method there can be an addition of ways to control for covariates in the CIERCE methods. For a more multivariate approach to Crosstalk, rather than a nested approach using sample pairs use combinations of samples to test for co-enrichment amongst the sample combination. This will require a brand-new statistical testing framework and thoughtful implementation to enable this enhancement to the CIERCE method.

In conclusion, the CIERCE methods are powerful approaches that empower researchers to explore, analyze, and connect sample-level gene signatures. By leveraging omics data and integrating Crosstalk and RECO, the CIERCE methods facilitate a deeper understanding of the landscape of multigene targets in cancer biology. It offers valuable insights into potential therapeutic targets and clinical implications. CIERCE has the potential to accelerate research in the field of cancer biomarkers and contribute to the development of more effective personalized treatment strategies for cancer patients.

## Methods

The CIERCE methods are implemented an R Package on GitHub for the analysis and visualization of gene signatures in cancer (https://github.com/kmlabdms/CIERCE). CIERCE provides researchers with a fast and convenient method for hypothesis generation focused gene signature analysis.

### EAS Derivation

CIERCE is populated by TCGA (patient data)^6^ and CCLE (patient-derived cell line data) data)^7^ to define sample-level EASs based on genes with (absolute) high expression and amplification.

TCGA RNA expression data was sourced from cBioPortal TCGA Pan-Can Atlas^16^. High expression is defined by within-gene across-sample RNA expression (microarray or RNA-Seq RPKM) z-scores relative to diploid samples provided by cBioportal of at least two. Amplification data is from UCSC Xena from TCGA Pan-Can (PANCAN) study’s gene level copy number defined by GISTIC 2.0 threshold ^17,18^. Amplification is defined based on putative copy number (CN) alterations from GISTIC2 ^17^ using an inferred (from copy number segment) log2 relative to diploid gene-level copy number of at least one. Each sample has their own EAS and that EAS may be defined by as few as a single gene. CCLE RNA expression data was sourced from cBioPortal ^16^. High expression is defined the same as TCGA. Amplification data is from cBioPortal gene level copy number defined by GISTIC 2.0 threshold. Amplification is also defined the same as TCGA. The populated omics data used for signature generation are obtained from cBioPortal ^16^ and USCS Xena ^18^.

### Pan-cancer EAS and Survival Analysis

The data used to create the EAS for this Pan-Can study were sourced from TCGA Pan-Can (PANCAN) study, encompassing 30 different cancer types and 8,650 samples. Forest plots were generated using the R package ggplot2 (v3.4.2)^19^. These plots were constructed based on hazard ratios and 95% confidence intervals obtained from Cox proportional hazard models that were fitted for Overall Survival (OS) ^20^ in relation to the binary prevalence, presence or absence, of EASs using R package survival (v3.5-7)^21^. Cancer types that violated the proportional hazards assumption were removed.

### Sample Phenotype and EASs

EAS Signatures are defined per sample, with the signature corresponding to either a TCGA sample ID or a CCLE cell line name for downstream exploration. Cancer types were defined by TCGA project for TCGA samples and by CCLE primary site for CCLE samples. Sample level molecular (e.g., FGA, TMB) and clinical (e.g., survival) features were obtained from cBioPortal ^16^. HER2+ status for TCGA samples was not included in the TCGA Pan Can Atlas on cBioPortal, but immunohistochemistry results for HER2, were available under TCGA Firehose Legacy for TCGA BRCA.

### Recurring Co-occurring (RECO) gene group

RECO genes are defined as highly prevalent genes that exhibit a co-occurrence of signature genes across samples. To extract RECO genes, high-prevalence genes are first identified, and unsupervised clustering using ComplexHeatmap (v2.16)^22,23^ is applied to detect their co-occurrence which is then checked using a clustering function and visualized using onco-plots.

### RECO gene group Significance Testing

Two bootstrapping methods can be run, and for a RECO gene group to be significant overall it had to be significant in both bootstrapping methods.

In the first method the number of prevalent samples for a given RECO group are randomly sampled with replacement from all samples used in the RECO analysis method. Then, within this random sample the number of samples prevalent for the RECO signature are counted. This is repeated 1,000 times, to generate a distribution of randomly prevalent samples for the RECO group’s prevalence in samples; p-values were calculated as the proportion of the distribution strictly greater than what was observed.

In the second method the number of genes for each RECO gene group is sampled from the distinct signature genes of all samples to create a random RECO gene set. Then, samples were tested for prevalence of this random RECO gene set. This is repeated 1,000 times to generate a distribution of sample prevalence for an equal length random RECO gene set; p-values were then calculated as the proportion of randomly generated prevalent samples strictly greater tan what was observed.

### Crosstalk

Crosstalk is designed to discover potential relations between signatures across systems using an adaptation of gene enrichment analysis based on the hypergeometric distribution. This process requires three gene sets: query gene sets, reference gene sets, and background gene sets (by default this is genome wide). For each sample-level signature pair constructed, a hypergeometric distribution is applied using the two defined gene sets and their significance recorded and visualized in a heatmap.

The output demonstrates relationships between sample pair signatures that are visualized using heatmaps and oncoplots^23^ of gene signatures, with the ability to be annotated with clinical and drug sensitivity data.

CIERCE also provides the ability to plot the resulting Crosstalk Matrix with 3D bars for each cell. This is best used with smaller sample sizes due to size of cells and thus bars, and it is best used within a cancer type and across model systems because they tend to have more variability in gene overlaps which is more informative in a 3D visualization.

When using Crosstalk with greater than 2 systems, we pose a nested approach. This Nested Crosstalk uses 2 systems as a baseline reference and query. To extend this to more systems, the sample pairs from the baseline Crosstalk are used as a reference for further query systems. Once the next Crosstalk is done the resulting Crosstalk matrix is collapsed via a union to be in reference to the baseline sample pairs. This can be done for as many query systems as provided. This approach avoids recursively adding sample pairs for each system and exponentially increasing computational time. This approach allows users to still get the power of Crosstalk without the burden of tremendous computational time. This approach is also superior to taking the union of the query systems and using only the baseline Crosstalk, as this finds co-enrichment across all systems, which won’t be tested if there is a union of the query systems.

Crosstalk can also be used for Enrichment Analysis as it finds significant pairs from reference and query. In CIERCE, RECO gene groups as query can be tested for enrichment in known signatures as reference to find more functional information about the given RECO gene group.

### Spatial Chromosome Locations

For a given set of RECO gene groups, gene locations were sourced from biomaRt^24,25^ v2.56.1. Gene locations were input into MG2C^26^ and SVG figures were output and edited for supplemental figures. Gene locations were put into terms of percent distance to telomere and centromere. For an arm of a chromosome (midpoint of centromere to start/end of chromosome), the percent distance to a centromere was defined as the absolute value difference between the gene location and the midpoint of the centromere, and the percent distance to a telomere was the distance to the closest either start or end of the chromosome. The sum of these two percent distances is 100% which is equal to the length of the arm of the chromosome the gene is on. These two distances were plotted for each case study and each RECO gene group.

### Co-expression networks

For a set of genes and a TCGA samples, the TCGA RNA expression data was subset to represent those samples and genes. This data was then input into GWENA ^27^ to find co-expression. Hub genes were defined as top 30% of connectivity for all genes.

## Acknowledgments

This work was supported by the University of Texas Dell Medical School Research Funds [to JK].

## Author Contributions

Initial manuscript draft: MY

Editing of manuscript: MY, QX, JK

Code Implementation: MY, QX

Analysis: MY, QX, JK

Conception: JK

## References

1 Rous, P. A SARCOMA OF THE FOWL TRANSMISSIBLE BY AN AGENT SEPARABLE FROM THE TUMOR CELLS. Journal of Experimental Medicine 13, 397–411 (1911). 10.1084/jem.13.4.397

2 Pivot, X. et al. Efficacy of HD201 vs Referent Trastuzumab in Patients With ERBB2-Positive Breast Cancer Treated in the Neoadjuvant Setting: A Multicenter Phase 3 Randomized Clinical Trial. JAMA Oncology 8, 698–705 (2022). 10.1001/jamaoncol.2021.8171

3 Bizzarri, M., Cucina, A., Conti, F. & D’Anselmi, F. Beyond the Oncogene Paradigm: Understanding Complexity in Cancerogenesis. Acta Biotheoretica 56, 173–196 (2008). 10.1007/s10441-008-9047-8

4 Chapman, P. B. et al. Improved Survival with Vemurafenib in Melanoma with BRAF V600E Mutation. New England Journal of Medicine 364, 2507–2516 (2011). 10.1056/NEJMoa1103782

5 Hahn, W. C. et al. An expanded universe of cancer targets. Cell 184, 1142–1155 (2021). 10.1016/j.cell.2021.02.020

6 Tomczak, K., Czerwińska, P. & Wiznerowicz, M. The Cancer Genome Atlas (TCGA): an immeasurable source of knowledge. Contemporary oncology 19, A68 (2015).

7 Ghandi, M. et al. Next-generation characterization of the Cancer Cell Line Encyclopedia. Nature 569, 503–508 (2019). 10.1038/s41586-019-1186-3

8 Taipale, J. et al. Effects of oncogenic mutations in Smoothened and Patched can be reversed by cyclopamine. Nature 406, 1005–1009 (2000). 10.1038/35023008

9 Walker, E. J. et al. Monoallelic Expression Determines Oncogenic Progression and Outcome in Benign and Malignant Brain Tumors. Cancer Research 72, 636–644 (2012). 10.1158/0008-5472.CAN-11-2266

10 Berger, A. C. et al. A Comprehensive Pan-Cancer Molecular Study of Gynecologic and Breast Cancers. Cancer Cell 33, 690–705.e699 (2018). 10.1016/j.ccell.2018.03.014

11 Karakas, B., Bachman, K. E. & Park, B. H. Mutation of the PIK3CA oncogene in human cancers. British Journal of Cancer 94, 455–459 (2006). 10.1038/sj.bjc.6602970

12 Zhu, G., Pei, L., Xia, H., Tang, Q. & Bi, F. Role of oncogenic KRAS in the prognosis, diagnosis and treatment of colorectal cancer. Molecular Cancer 20 (2021). 10.1186/s12943-021-01441-4

13 Underhill, M. L., Germansky, K. A. & Yurgelun, M. B. Advances in Hereditary Colorectal and Pancreatic Cancers. Clinical Therapeutics 38, 1600–1621 (2016). 10.1016/j.clinthera.2016.03.017

14 Wang, S. et al. The molecular biology of pancreatic adenocarcinoma: translational challenges and clinical perspectives. Signal Transduction and Targeted Therapy 6 (2021). 10.1038/s41392-021-00659-4

15 Tipney, H. & Hunter, L. An introduction to effective use of enrichment analysis software. Human Genomics 4, 202 (2010). 10.1186/1479-7364-4-3-202

16 Cerami, E. et al. The cBio cancer genomics portal: an open platform for exploring multidimensional cancer genomics data. Cancer Discov 2, 401–404 (2012). 10.1158/2159-8290.CD-12-0095

17 Mermel, C. H., Schumacher, S. E., Hill, B., Meyerson, M. L., Beroukhim, R. & Getz, G. GISTIC2.0 facilitates sensitive and confident localization of the targets of focal somatic copy-number alteration in human cancers. Genome Biology 12, R41 (2011). 10.1186/gb-2011-12-4-r41

18 Goldman, M. J. et al. Visualizing and interpreting cancer genomics data via the Xena platform. Nature Biotechnology 38, 675–678 (2020). 10.1038/s41587-020-0546-8

19 Wickham, H. ggplot2: Elegant Graphics for Data Analysis. Springer-Verlag New York (2016). 10.1007/978-3-319-24277-4

20 Liu, J. et al. An Integrated TCGA Pan-Cancer Clinical Data Resource to Drive High-Quality Survival Outcome Analytics. Cell 173, 400–416.e411 (2018). 10.1016/j.cell.2018.02.052

21 T, T. A Package for Survival Analysis in R. R package version 3.5-7 (2023).

22 Gu, Z. Complex heatmap visualization. iMeta e43 (2022). 10.1002/imt2.43

23 Gu, Z., Eils, R. & Schlesner, M. Complex heatmaps reveal patterns and correlations in multidimensional genomic data. Bioinformatics 32, 2847–2849 (2016). 10.1093/bioinformatics/btw313

24 Durinck, S. et al. BioMart and Bioconductor: a powerful link between biological databases and microarray data analysis. Bioinformatics 21, 3439–3440 (2005). 10.1093/bioinformatics/bti525

25 Durinck, S., Spellman, P. T., Birney, E. & Huber, W. Mapping identifiers for the integration of genomic datasets with the R/Bioconductor package biomaRt. Nature Protocols 4, 1184–1191 (2009). 10.1038/nprot.2009.97

26 Chao, J. et al. MG2C: a user-friendly online tool for drawing genetic maps. Molecular Horticulture 1, 16 (2021). 10.1186/s43897-021-00020-x

27 Lemoine, G. G., Scott-Boyer, M.-P., Ambroise, B., Périn, O. & Droit, A. GWENA: gene co-expression networks analysis and extended modules characterization in a single Bioconductor package. BMC Bioinformatics 22, 267 (2021). 10.1186/s12859-021-04179-4

