## Supplemental Figures for "Frequently Together, Among Model Systems Molecular Alterations: Cancer Maintenance Markers with Translational Study Design"

Supplemental Figure 1. **Sample-, gene-level expressed amplification signatures (EAS) are highly prevalent among solid tumors based on a TCGA Pan-Can analyses.**

**a**, The distribution of the number of samples with EASs by TCGA Pan-Can project.

**b**, A boxplot distribution of the number of genes represented among EAS-containing samples with enumerated medians by TCGA Pan-Can project.

**c**, Forest plot of hazard ratios and 95% confidence Intervals comparing overall survival (OS) between samples with versus without (as the referent) EASs by TCGA Pan-Can project. Cancer types that violated the proportionality assumption were removed. Samples with EAS's that are associated with significantly ($p \le 0.05$) shorter- (in red) and longer-survival (in blue) are highlighted.

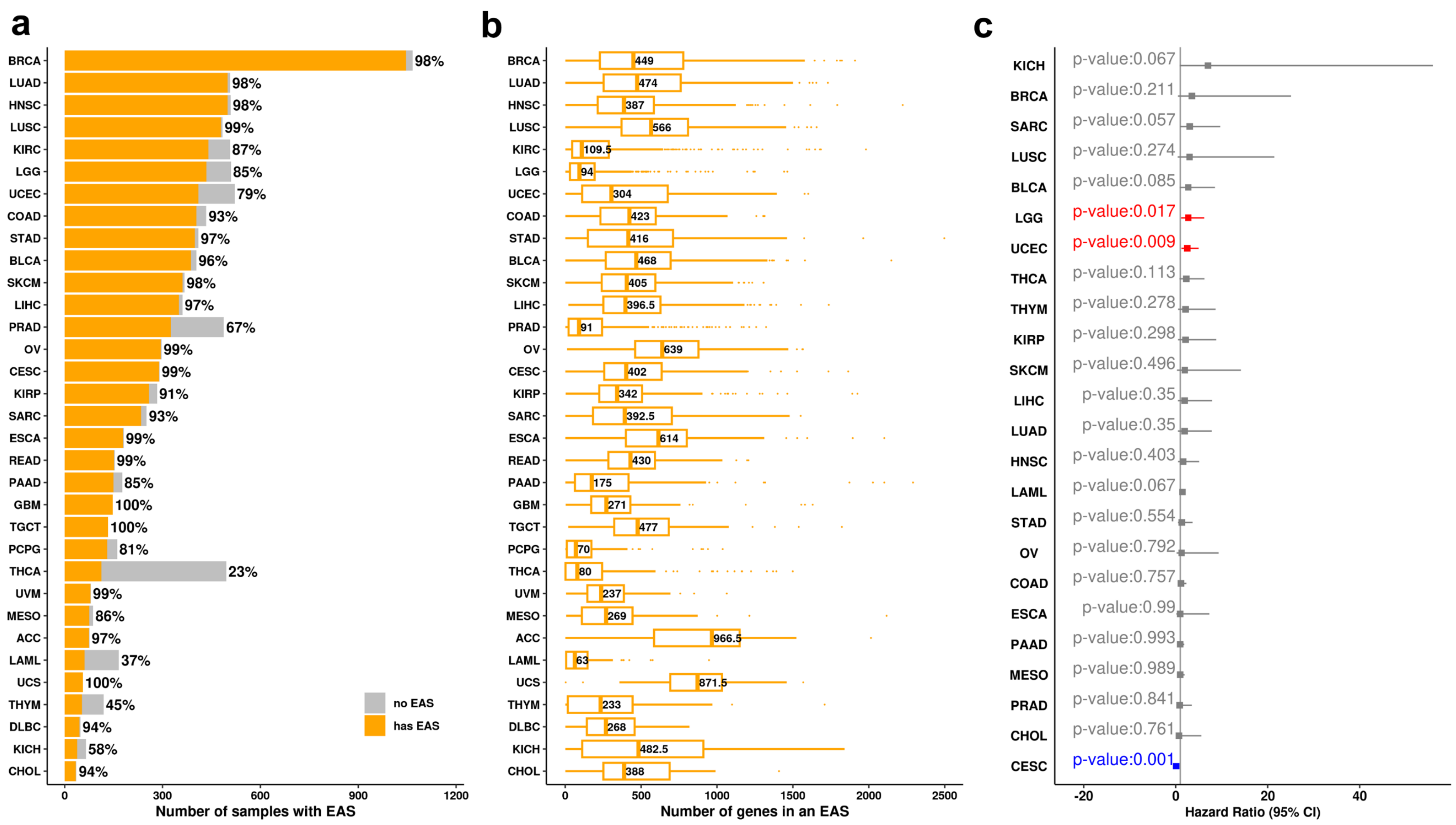

Supplemental Figure 2. **Sample-, gene-level expressed amplification signatures (EAS) are highly prevalent among solid tumors patient-derived cell lines based on a CCLE Pan-Can analyses.**

**a**, The distribution of the number of samples with EASs by CCLE Pan-Can project.

**b**, A boxplot distribution of the number of genes represented among EAS-containing samples with enumerated medians by CCLE Pan-Can project.

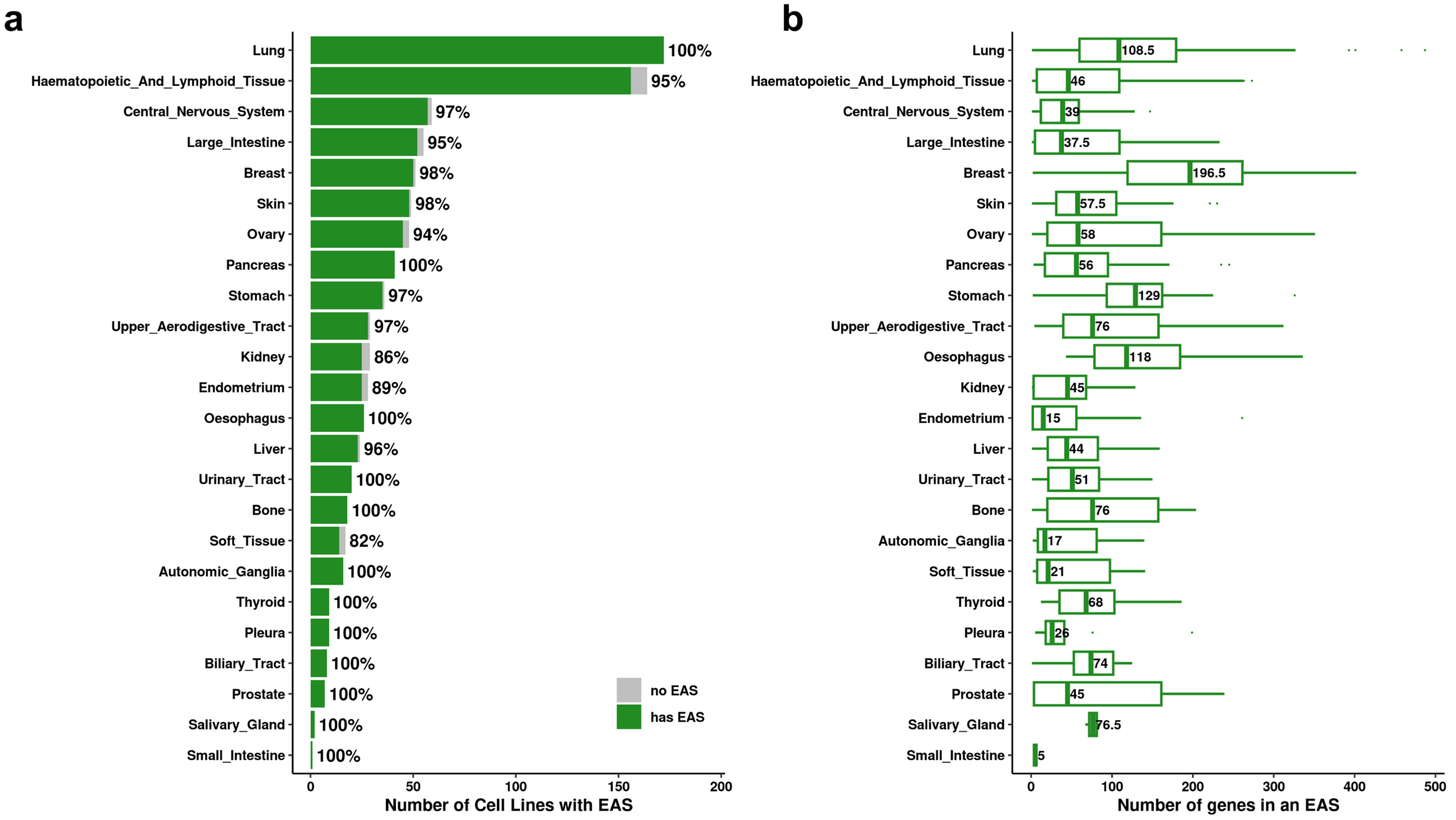

Supplemental Figure 3. **A RECO analysis of TCGA LGG samples reveals spatial heterogeneity in non-oncogene containing RECO gene groups, except for one, within Chromosome 7, each with varying co-expression networks.**

**a**, The mapping of RECO genes onto chromosome locations colored by RECO EAS gene group. LUZP6 was omitted as no location was present in biomaRt for the current build.

**b,** RECO EAS gene group co-expression networks. Hub genes (top 30% connectivity) are in yellow; non-hub genes are in purple. Line width corresponds to the level of connectivity. Networks with a level of connectivity (indicated by line width) of at least 0.9999 are shown.

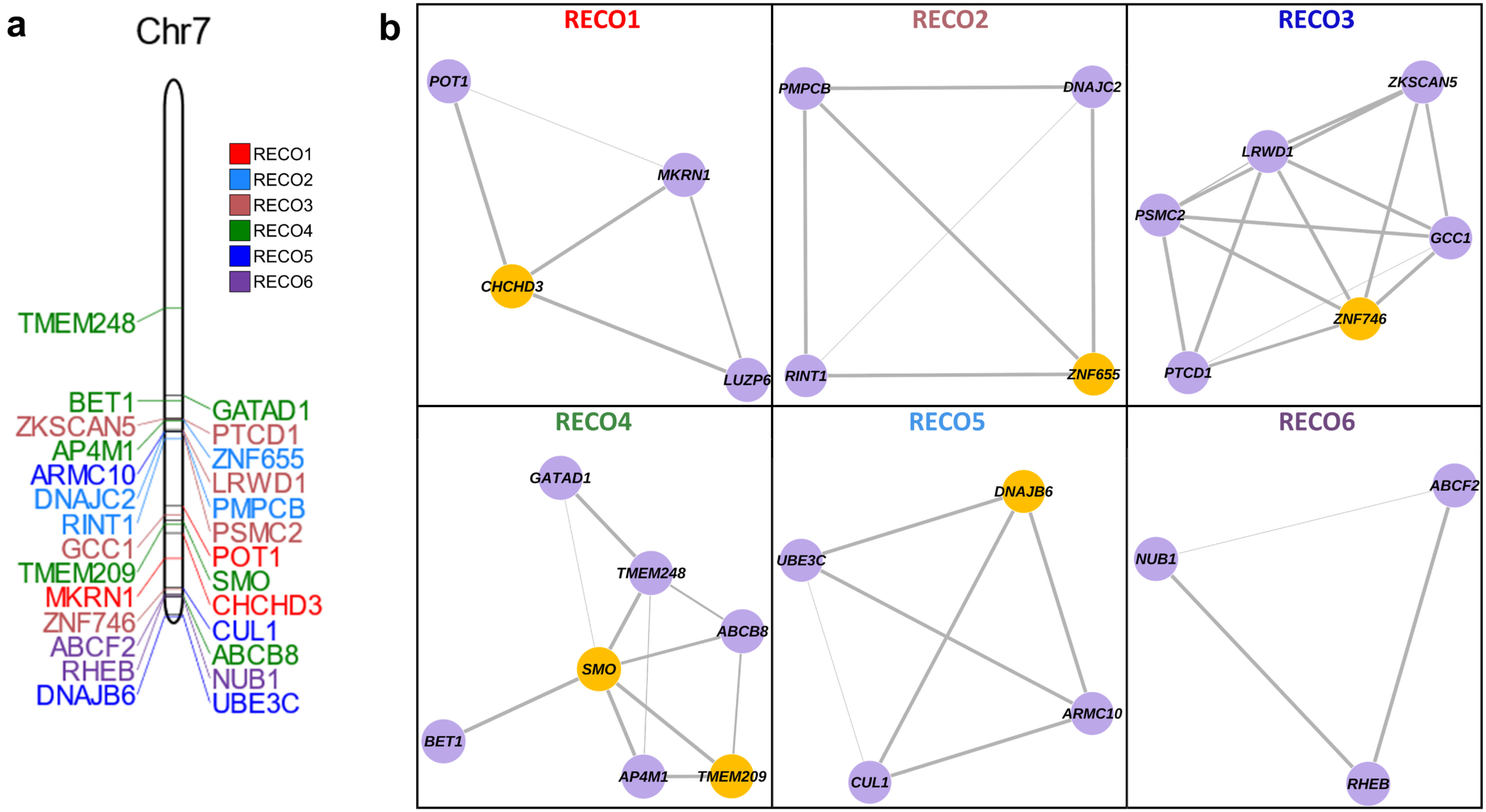

Supplementary Figure 4. **An assessment of (signature-containing) gene-level prevalence between query and reference model systems based on an analysis of residuals from equality between them detects differential over-representation.**

**a,** Case 2. For a given gene contained within a signature, the number of samples containing that gene in the query (TCGA BRCA HER2+) as compared to the reference (CCLE BRCA HER2+) by computing the residual from the equality (y=x) line for that gene shows a greater prevalence in TCGA.

**b,** Case 3. For a given gene contained within a signature, the number of samples containing that gene in the baseline query (TCGA UCEC) as compared to the reference (TCGA OV) by computing the residual from the equality (y=x) line for that gene shows an equal prevalence between query and reference systems.

**c,** Case 4. For a given gene contained within a signature, the number of samples containing that gene in the baseline query (TCGA CRC) as compared to the reference (TCGA Pancreatic) by computing the residual from the equality (y=x) line for that gene shows an equal prevalence between query and reference systems.

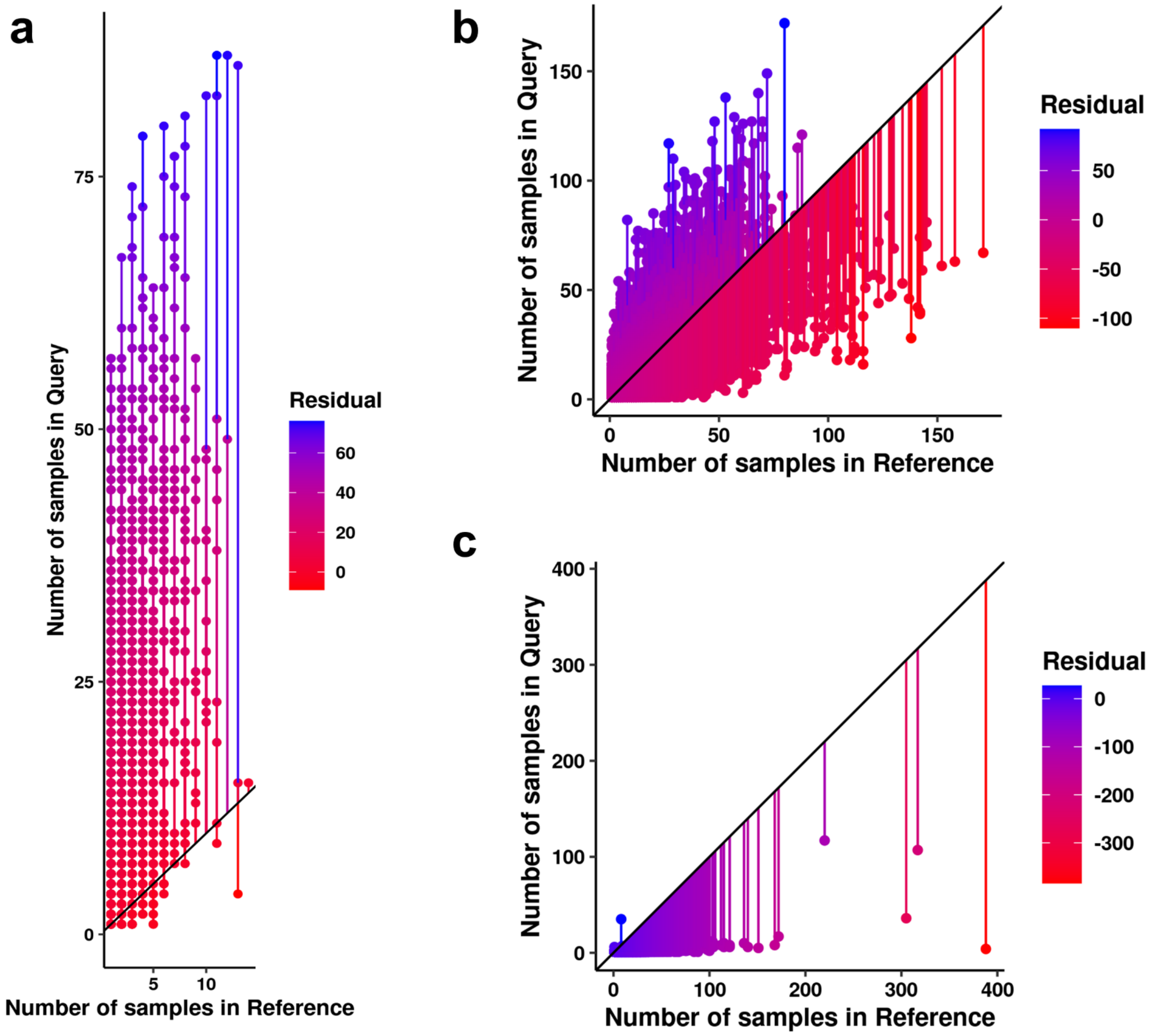

Supplemental Figure 5. **A crosstalk and RECO analysis of HER2+ breast cancer TCGA patients and cell lines EAS's reveals distinct spatial associations among non-oncogene containing RECO gene groups, except for one, within chromosomes 1, 8, 17, and 20 that show a high co-expression connectivity residing on Chromosome 8.**

**a**, The mapping of RECO genes onto chromosome locations.

**b,** RECO EAS gene group co-expression networks. Hub genes (top 30% connectivity) are in yellow; non-hub genes are in purple. Line width corresponds to the level of connectivity. Networks with a level of connectivity (indicated by line width) of at least 0.90 (RECO1-6) and 0.26 (RECO8) are shown.

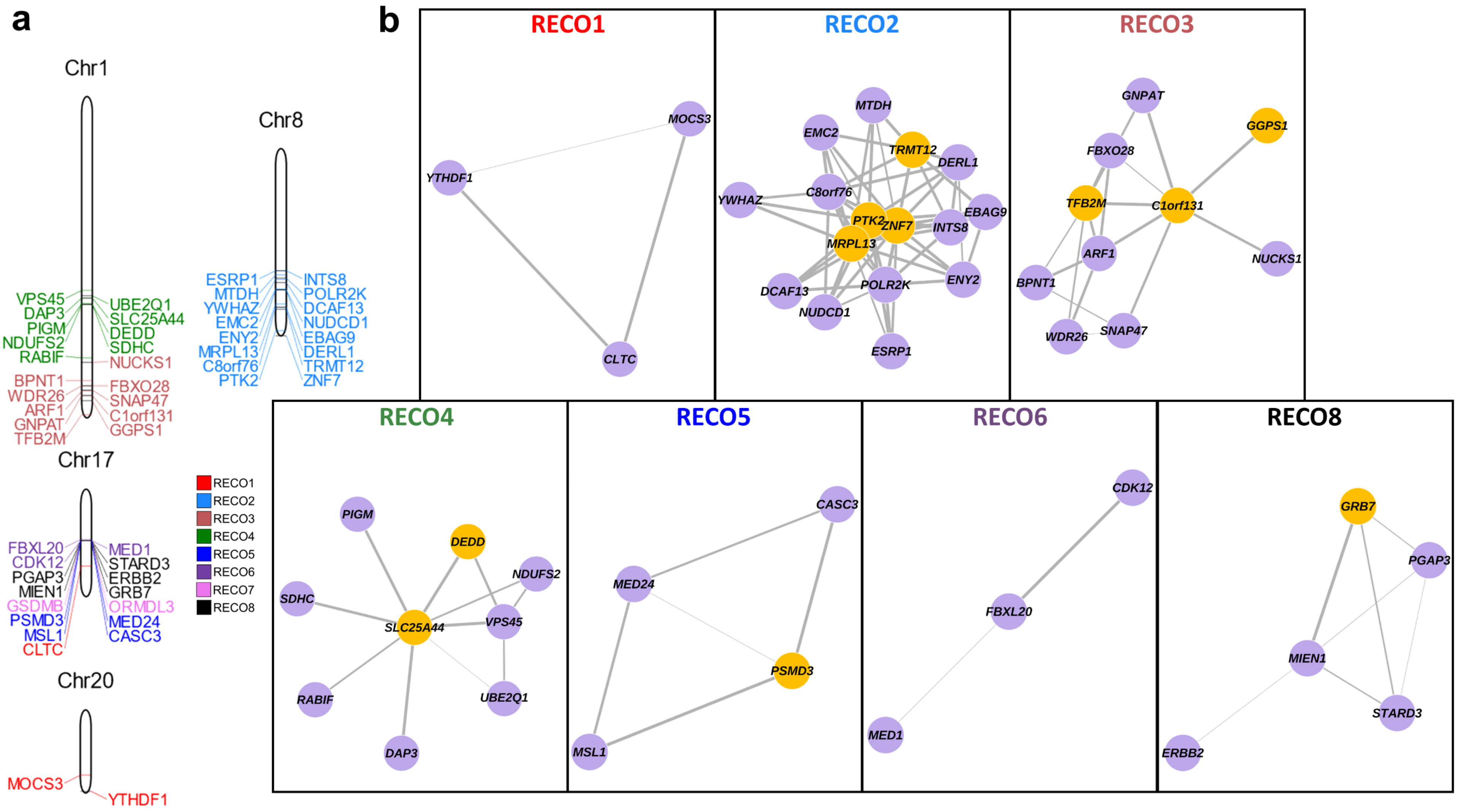

Supplemental Figure 6. **A nested crosstalk and RECO analysis of TCGA Pan-Gyn-Can EAS's reveals spatial heterogeneity in non-oncogene containing RECO gene groups, except for one, within each of chromosomes 1, 3, and 8 in which at least two RECO gene groups are defined, each with varying co-expression levels.**
**a**, The mapping of RECO genes onto chromosome locations.
**b**, RECO EAS gene group co-expression networks. Hub genes (top 30% connectivity) are in yellow; non-hub genes are in purple. Line width corresponds to the level of connectivity. Networks with a level of connectivity (indicated by line width) of at least 0.99 are shown.

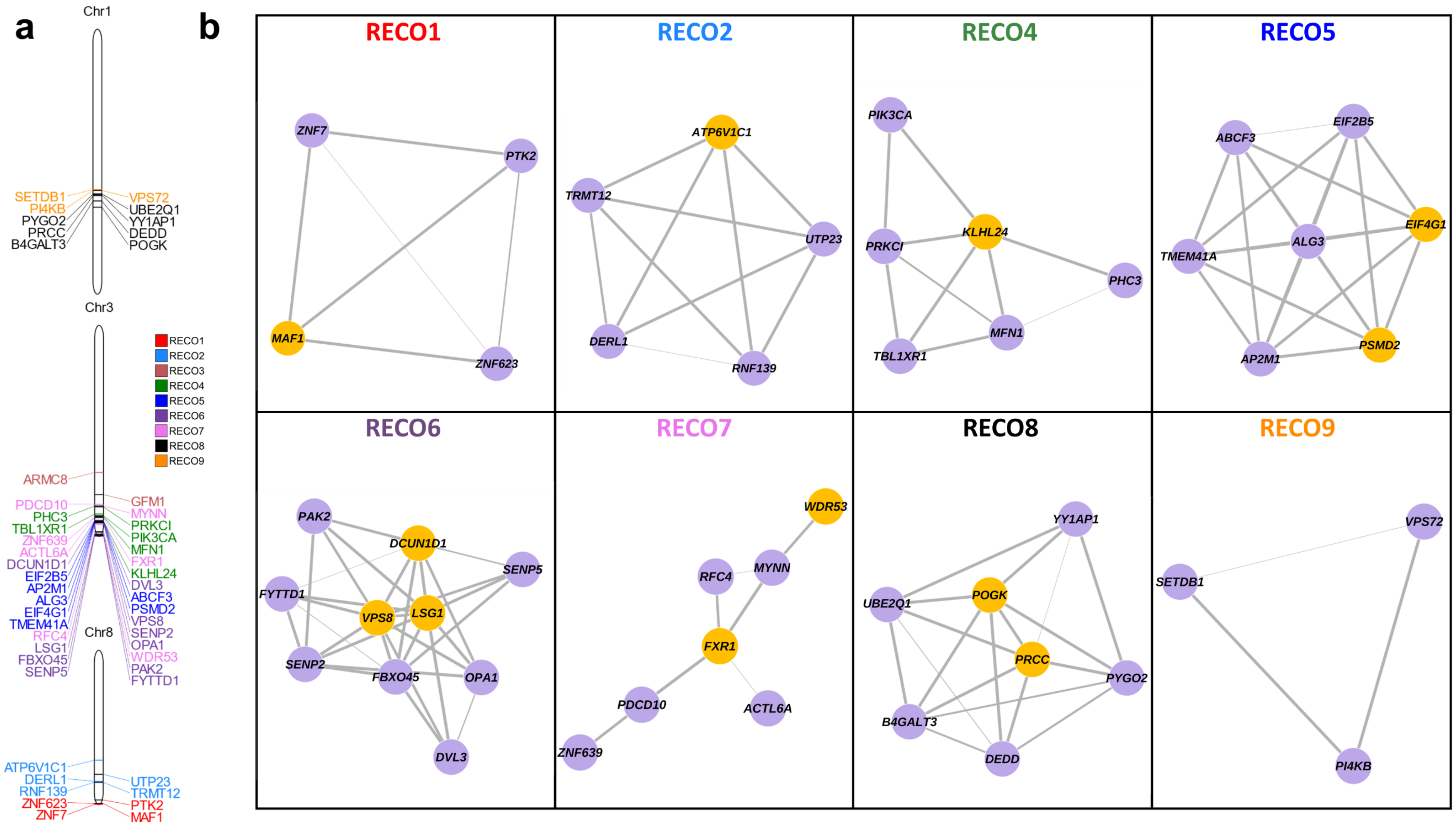

Supplemental Figure 7. **A Nested Crosstalk and RECO analysis of TCGA colorectal and pancreatic, and DNA damage repair (DDR) gene signature shows lack of a chromosome spatial association, though co-expression among RECO gene groups.**

**a,** The mapping of RECO genes onto chromosome locations

**b**, RECO EAS gene group co-expression networks. Hub genes (top 30% connectivity) are in yellow; non-hub genes are in purple. Line width corresponds to the level of connectivity. Networks with a level of connectivity (indicated by line width) of at least 0.99 are shown.

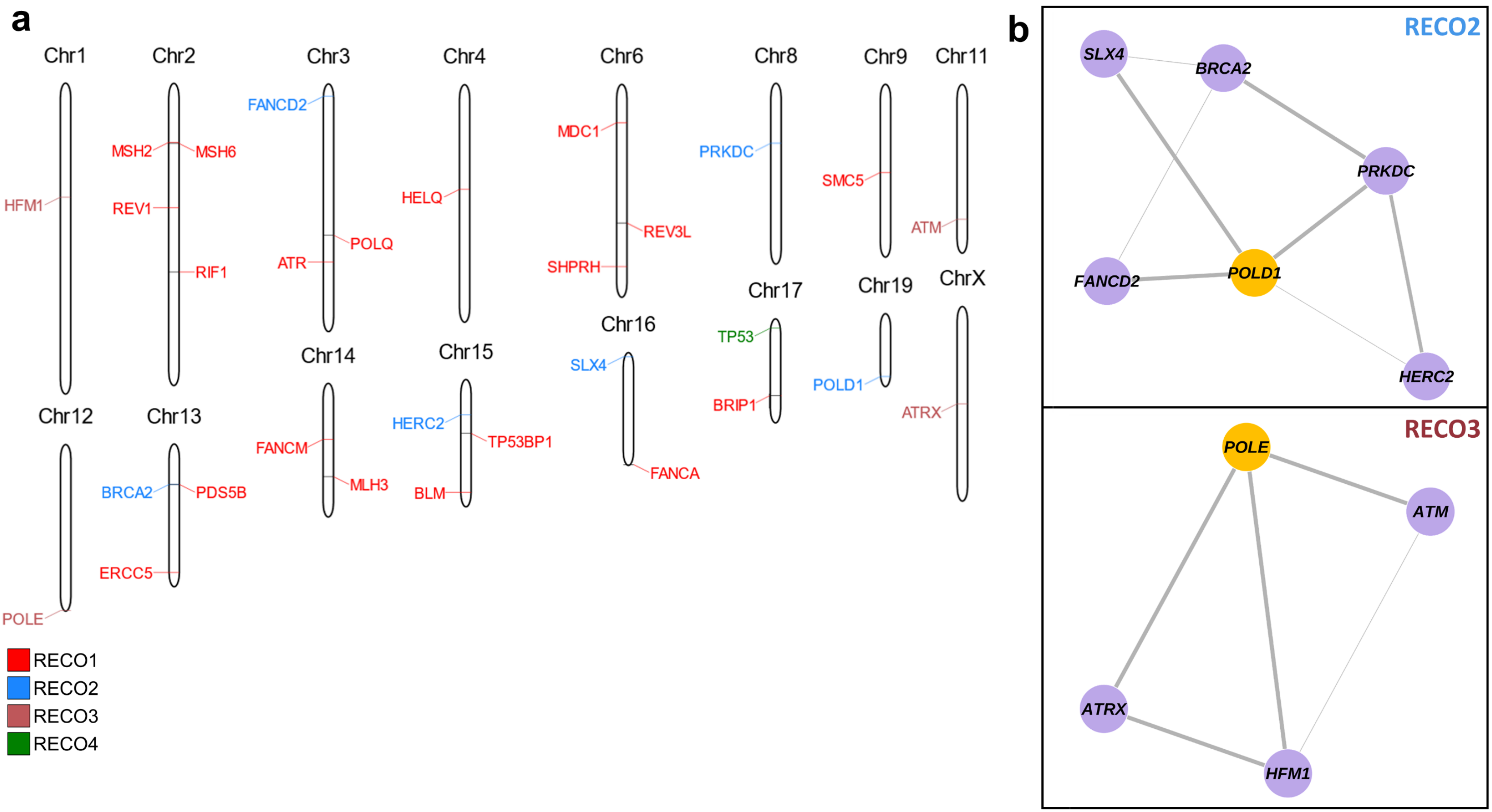

Supplemental Figure 8. **RECO gene groups depict differential spatial clustering relative to centromere and telomere locations.** Plot rows are case studies, and plot columns are the RECO gene groups; case studies without representative RECO gene groups are empty. Vertical and horizontal lines in red are provided to assess 50% (the center of the arm of the chromosome) in either direction towards centromere or telomere. Each point is a gene, and each gene will lie on the black line as its position a linear combination of relational locations to centromere and telomere. Genes in the bottom right corner of the plot will be closer to the centromere, and genes in the top left corner will be closer to the telomere.

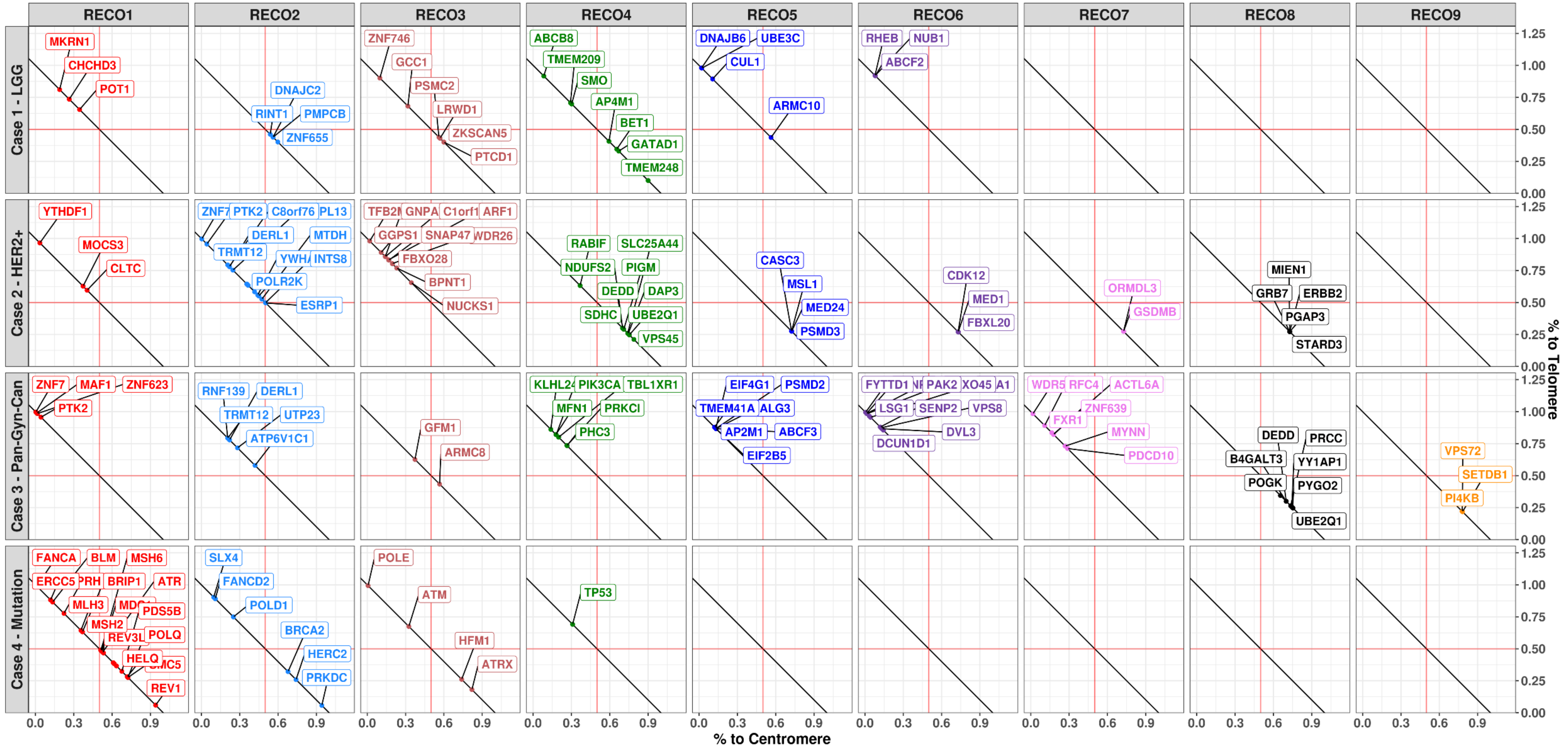
